# Experimental Thermal Stress Increases Corallicolid Relative Abundance in the Stony Coral *Pocillopora damicornis*

**DOI:** 10.64898/2026.08.17.745262

**Authors:** Hailey G. Znamenacek, Ella R. Wilson, Anthony M. Bonacolta, Kirsten S. Brendtro

## Abstract

Rising ocean temperatures disrupt previously stable coral-microbe interactions, leading to widespread coral mortality and threatening reef ecosystems worldwide. Growing evidence demonstrates the coral microbiome, including protists, plays a critical role in the host response to thermal stress. Specifically, corallicolids (Phylum: Apicomplexa) are positively correlated with thermal stress mortality in soft corals. This study investigates changes in the eukaryotic microbiome of the stony coral, *Pocillopora damicornis,* across an experimental thermal stress event. Using anti-metazoan 18S rRNA gene metabarcoding, protist communities were assessed at four time-points during experimental thermal stress. Outside of the Symbiodiniaceae, a prominent shift in microbiome composition during thermal stress was observed, most notably a significant increase and dominance in Corallicolida abundance in heat-stressed corals, while other protists declined substantially. Increased corallicolid abundance concurrent with bleaching suggests an overlooked compounding stressor beyond the loss of algal symbionts during heat stress. These results contrast with previous research on *Pocillopora* microbiomes showing prokaryotic community stability throughout stress, and support the hypothesis that thermal stress may alter the coral-corallicolid relationship, potentially shifting corallicolids from a commensal to a parasitic role, and synergistically contributing to coral mortality during and after heat stress. This work provides critical insight into the role of protists in marine holobionts, supports their inclusion in future microbiome studies, and informs strategies to improve coral resilience under climate change.

## INTRODUCTION

Coral reefs are among the most biologically diverse and ecologically important ecosystems on Earth, yet they are increasingly threatened by climate change-driven ocean warming. Rising ocean temperatures have triggered widespread coral bleaching, leading to a dramatic increase in coral mortality across the globe and the overall degradation of reef ecosystems (Hughes et al. 2018). While coral bleaching is characterized as the loss of symbiotic dinoflagellates (Family: Symbiodiniaceae; also commonly referred to as zooxanthellae), growing evidence suggests that other members of the coral microbiome also significantly impact host responses to thermal stress (Boilard et al. 2020). The bacterial portion of the coral microbiome has been extensively studied and shown to play a central role in regulating coral stress responses, with shifts in bacterial community composition closely linked to thermal stress, bleaching events, and overall coral resilience (Voolstra et al. 2024). The eukaryotic portion of the coral microbiome has been comparatively much less studied beyond the Symbiodiniaceae, despite the potential role of protists such as green algae and apicomplexans to overall coral host health and fitness (Bonacolta et al. 2023). Corallicolids, for instance, are apicomplexans commonly found associated with anthozoans across the globe, including *Pocillopora damicornis* (Clerissi et al. 2018; Kwong et al. 2019). Corallicolids appear to have an intimate, context-dependent symbiotic relationship with their coral host. Under favorable conditions, corallicolids appear to be commensal, as no known negative impact on coral host health has been observed under normal conditions (Keeling et al. 2021). However, recent studies have shown that corallicolids are found in higher abundance in thermally susceptible soft corals and even frequently increase in abundance during natural heat-stress events (Bonacolta et al. 2024; Peterson et al. 2025), suggesting that environmental stress may alter their role within the coral host.

The genus *Pocillopora* comprises 15 species, and dominates reef ecosystems as key reef builders providing marine habitat for many invertebrates and commercially important fish species. *P. damicornis,* also known as the cauliflower coral, is native to the tropic and subtropic regions of the Indian and Pacific Oceans, where it is one of the most prevalent coral species (Veron et al. 2025). *P. damicornis* is generally considered moderately resistant to environmental stressors relative to other coral species. Rapid temperature spikes or cold shocks can still trigger bleaching; however, its prokaryotic microbiome typically remains stable (Bergman et al. 2021). Although often overlooked in microbiome studies, this species possesses a diverse, protist-rich microbiome, including consistently high abundances of corallicolids (Zallio et al. 2026), making it an excellent model for investigating coral host-associated protist responses to thermal stress. Previous metabarcoding studies revealed that corallicolids are one of the most abundant and widespread non-Symbiodiniaceae protists associated with *P. damicornis* across the globe (Clerissi et al. 2018; del Campo et al. 2026; Zallio et al. 2026). In addition to corallicolids, MALVs (marine alveolates), a diverse group of parasitic dinoflagellates, are often found within coral-associated protist communities and may contribute substantially to microeukaryotic diversity (Clerissi et al. 2018; Bonacolta et al. 2023). The presence of these protists across diverse environmental conditions suggests they are consistent associates of the coral microbiome and may play important roles in host health and stress responses.

Thermal stress disrupts the sensitive balance between corals and their symbiotic microalgae by impairing photosynthesis and triggering the expulsion of Symbiodiniaceae, leading to bleaching (Vidal-Dupiol et al. 2009). While much of the bleaching response is rightfully attributed to this algae loss, increasing evidence suggests that microbial community shifts may accelerate this process and increase the likelihood of coral mortality following bleaching (Gardner et al. 2019). Corallicolids, in particular, have been positively associated with thermally susceptible soft corals, raising the possibility that they contribute to microbiome instability under stress, in contrast to other protists in the coral microbiome, such as MALVs, which are positively associated with thermally resilient soft corals (Bonacolta et al. 2024). However, what drives this shift, and the extent to which corallicolids induce it, remains poorly understood.

This study investigates changes in the protist community of *P. damicornis* across experimental thermal stress. Using anti-metazoan 18S rRNA gene metabarcoding, protist abundance and community dynamics are examined through experimental thermal stress. By examining coral-associated protist community responses to thermal stress, this work aims to clarify the role of protists in coral microbiome instability and bleaching response. Improved understanding of symbiotic protist behavior under heat stress may provide critical information on coral vulnerability in warming oceans and help future efforts to predict and mitigate coral bleaching under climate change.

## MATERIALS AND METHODS

Ten fragments (∼9-20 cm) of the coral *P. damicornis* were maintained in a custom aquarium under controlled oceanic conditions (salinity 35 ppt; temperature 20°C) and sampled throughout the experiment (see Supplementary Figure 1). Corals were sourced from Underwater Wonders (Fort Collins, Colorado). Chemistry conditions were maintained for magnesium (1250-1400 ppm), calcium (380-450 ppm), nitrate (5-10 ppm), phosphorus (0.02-0.10 ppm), pH (∼8.2), and alkalinity (8-12 dKH). Levels of each condition were monitored and maintained within optimal ranges. Magnesium, nitrate, and calcium levels were tested using the Salifert Profi Test. Phosphorus and alkalinity were tested using a Hanna Instruments handheld colorimeter (Hanna Instruments Inc, Woonsocket, RI, USA). The EnviroGroⓇ T5 high-output fluorescent 120 V light (Petaluma, California) provided illumination for the tank, mimicking a diurnal light cycle.

The health of the corals was monitored through visual observation and standardized photographic documentation of specific fragments (Supplementary Figure 2). A grid system within the tank allowed for random selection of the coral when sampling, ensuring an unbiased representation across varying coral fragment health states. Once the environmental conditions stabilized, six samples were taken, using the randomized selection process. Sampling was performed using autoclaved forceps (one per sample), and the coral tissue was transferred into labeled 2mL microcentrifuge tubes corresponding to grid positions (See Supplementary Figure 1 for a schematic of the experimental design).

Sampling was conducted across four experimental conditions: Control (C0; 20°C, week 5), Mild Stress (MS; 22.2°C, week 8), Cold Stress (CS; 18°C, week 9), and Heat Stress (HS; 27.2°C, week 15) (Supplementary Figure 1). Temperature increase was achieved using a submersible heating probe. MS samples were collected as the corals started exhibiting paling (22.2°C, week 8), and HS samples were collected once the corals became translucent and showed reduced polyp-extension. The CS group was not in the original experimental plans, but occurred after an attempt to slow the rapid increase in tank temperature; polyp samples were taken one week after the MS group at 18°C.

Genomic DNA was extracted and prepared using the ZYMO RESEARCH *Quick*-DNA MiniPrep Kit (Solid Tissue protocol; Irvine, California, USA). A previously validated nested PCR approach was implemented to decrease the host signal for protist microbiome studies (del Campo et al. 2019). First, PCR pre-amplification of non-metazoan eukaryotic 18S rRNA genes was conducted using primers 18S-EUK581-F (5’-GTGCCAGCAGCCGCG-3’) and 18S-EUK1134-R (5’-TTTAAGTTTCAGCCTTGCG-3’), each diluted 10uM prior to use. Each 25 uL reaction contained 12.5 uL of 2x Phusion Flash PCR Master Mix, 1uL of each primer, 7.5uL UltraPure Water, and 3uL of DNA. Thermal cycling conditions for PCR consisted of: 30 seconds at 98°C. 35 cycles of 10 seconds at 98°C, 30 seconds at 51.1°C, and 1 min at 72°C. The final extension was 5 min at 72°C. Once completed, amplicons were refrigerated at 4℃. Amplification success was determined by agarose gel electrophoresis, and successfully amplified samples were selected for sequencing. After running gels, the weakest sample from each group was removed, discovered through gel band intensity. Five samples were selected for each timepoint (C0, MS, CS, and HS). Samples were prepared according to Novogene preshipment protocols (Novogene-Preshipping Checklist, and the Sample Information Form) and submitted for further amplification and sequencing using the following forward and reverse primers: E572-F (5’-CYGCG GTAAT TCCAG CTC-3’) and E1009-R (5’-AYGGT ATCTR ATCRT CTTYG-3’). These primers for this second round of PCR fit within the previously amplified non-metazoan amplicons from the first round of PCR (del Campo et al. 2019). Sequencing was performed with a target depth of 100,000 reads per sample on an Illumina Novaseq 6000.

Sequencing data was analyzed in R Studio (R version 2025.05.1+513) using standard microbiome analysis workflows and protocols described in (Bonacolta et al. 2024), following trimming of primers and sequencing adaptors with Cutadapt (Martin 2011). Trimmed sequences were processed using the DADA2 pipeline (Callahan et al. 2016), with forward and reverse reads truncated at 225 bp and 210 bp, respectively, based on quality score profiles prior to read merging. Amplicon sequence variants (ASVs) were generated and taxonomically classified using the PR2 Database (Guillou et al. 2013). Shannon alpha diversity indices were calculated to evaluate within-sample diversity, while beta diversity was assessed using Aitchison distances to compare community structure between the different stress conditions. A Global Wilcoxon Test with an adjusted p-value calculated using the “Holm” pairwise comparison was conducted to determine whether alpha diversity differed significantly between sample points, while beta diversity was statistically tested using ANOSIM within the vegan R package (Oksanen et al. 2025). A bubble plot was created to visualize the taxonomic make-up of the protist communities through heat-stress using Phyloseq (McMurdie and Holmes 2013), ggplot2 (Wickham et al. 2019), and tidyverse tools. Analysis of Compositions of Microbiomes with Bias Correction (ANCOM-BC) tests were run to statistically test for significance in microbial abundance changes due to thermal stress (Lin and Peddada 2020).

## RESULTS

The eukaryome of CS *P. damicornis* exhibited significantly higher Shannon Alpha Diversity relative to C0 and HS samples, with shifts in abundance evident across multiple taxa (Figure 1A, Figure 2). MS samples showed significantly higher Shannon Alpha Diversity than C0 (Figure 1A). This elevated diversity likely reflects a dysbiosis under thermal stress, potentially driven by an increase in rarer or opportunistic taxa. Corallicolida (Apicomplexa) were present at moderate levels at this stage. Other protist taxa remained detectable in MS samples, indicating a partial community shift rather than dominance by a single taxon. By the HS timepoint, however, protist community composition shifted markedly: corallicolids became dominant, while many other protist genera declined to very low abundance or fell below detection (mean relative abundance <0.1%, excluding Symbiodiniaceae; Figure 2). This shift toward a single dominant taxon corresponded with a significant decline in Shannon Alpha Diversity from MS to HS (Wilcoxon, P=0.016; Figure 1A), reflecting this transition toward an apicomplexan-dominated eukaryome.

**FIGURE 1.**
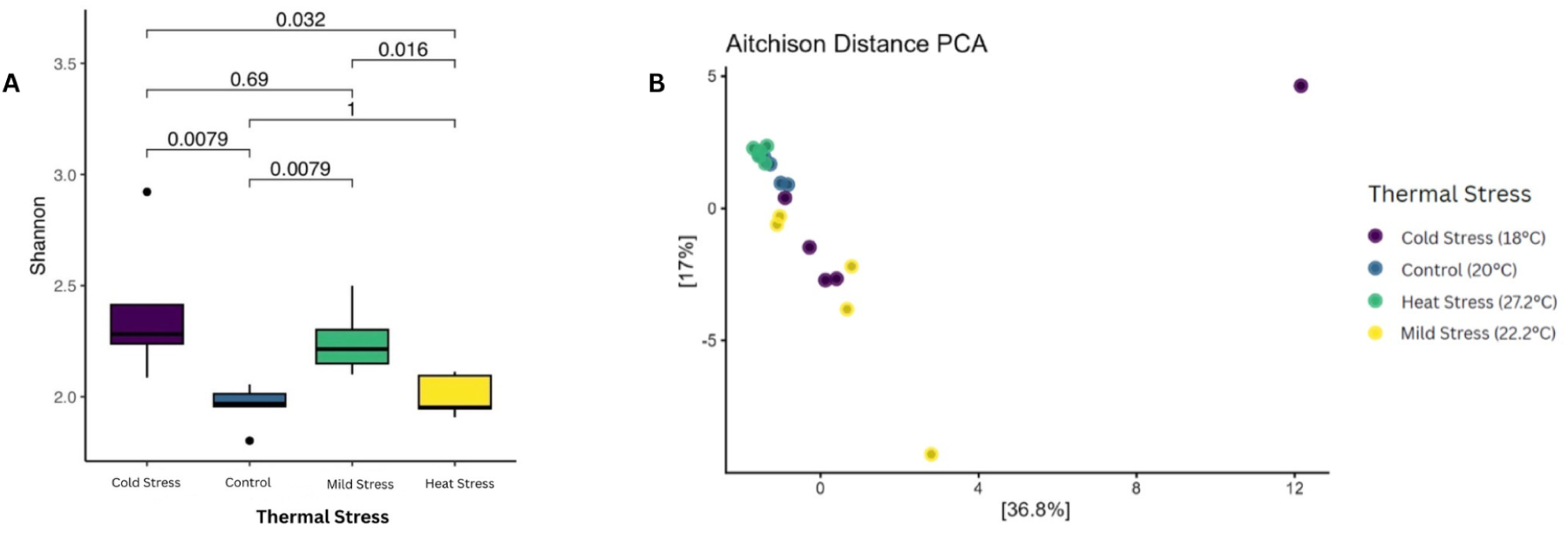
*Pocillopora damicornis* community indices through thermal stress. **(A) Alpha Diversity Plot.** Shannon diversity was used to assess within sample-protist diversity. Each block represents a different stress event, and the p-values indicate the statistical comparisons between each stress event. A Global Wilcoxon Test with an adjusted p-value calculated using the “Holm” pairwise comparison was used to determine significance. Cold and mild stressed corals exhibited significantly greater alpha diversity than controls (p= 0.0079), while heat stressed corals had significantly lower diversity than both cold (p= 0.032) and mild stressed corals (p= 0.016). **(B) Aitchison Distance Beta Diversity PCA.** Control and heat stressed samples each form relatively tight clusters, indicating consistent protist community composition between treatments. In contrast, mild and cold stress samples are more dispersed, reflecting greater variability in the protist community during transitional or abrupt stress conditions creating a destabilized microbiome. Distinct clustering patterns among thermal stress conditions are shown, the cold stress group includes a strong outlier, suggesting an extreme community shift in at least one cold shocked coral. Under control conditions (C0), the protist community is relatively diverse and evenly distributed across multiple taxonomic groups. Coloring is based on the thermal conditions throughout the experiment.

**FIGURE 2.**
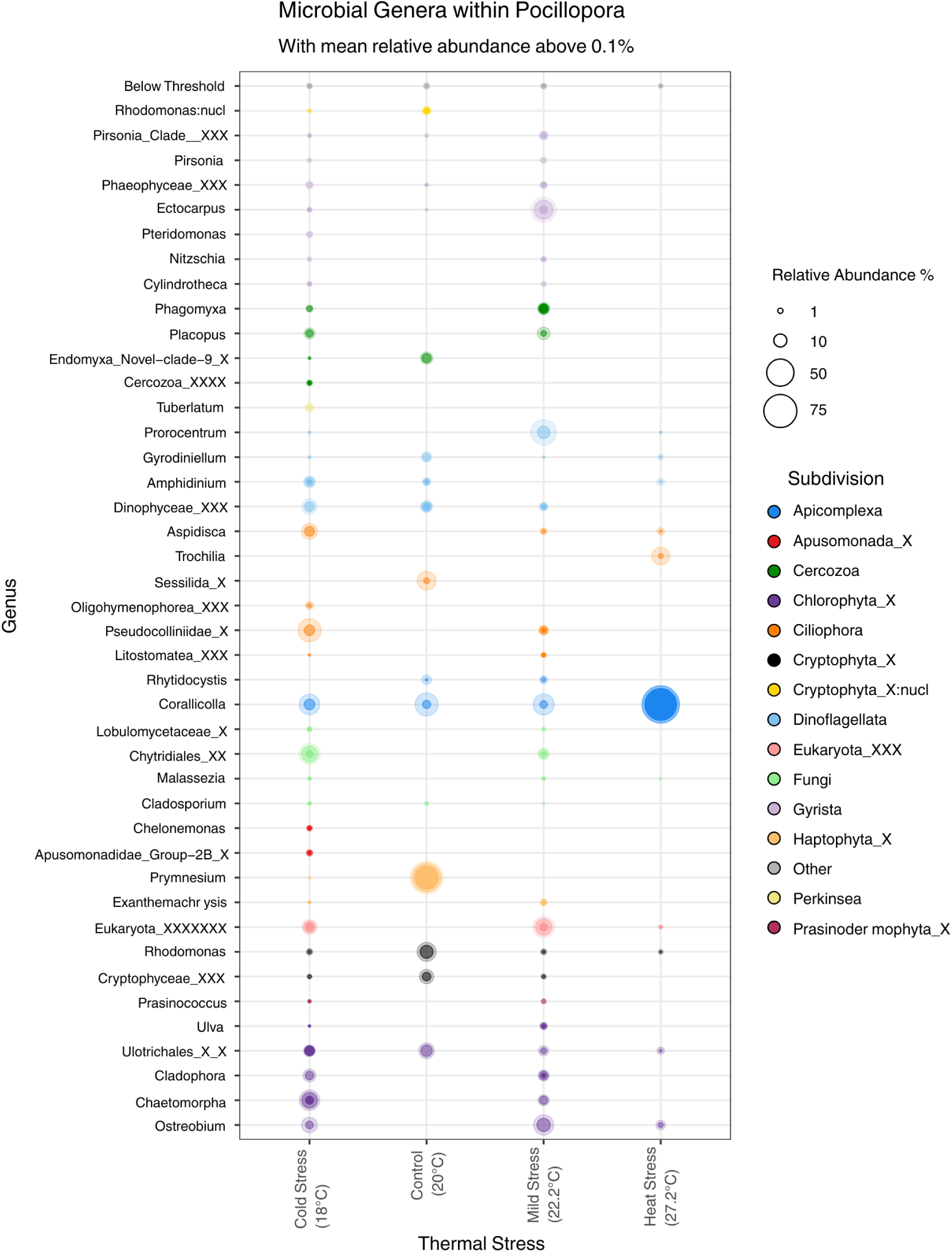
Bubble plot of protist genera associated with *Pocillopora damicornis* across four thermal conditions: cold stress (CS), control (C0), mild thermal stress (MS), and heat stress (HS). Only genera with a median relative abundance above 0.01% are shown, allowing comparison of the predominant taxa while excluding insignificant groups. Symbiodiniaceae were excluded from the chart due to their prominence in the microbiome (See also Supplementary Figure 3). Lower confidence taxonomic labels were given unique suffixes according to their highest assigned taxonomic rank (e.g., _X, _X_X, _XXX).

Protist composition within *P. damicornis* differed significantly across thermal stress treatments (C0, MS, CS, HS; ANOSIM, R=0.273, P=0.001), though the moderate R value indicates partial rather than complete separation between groups (Figure 1B). Symbiodiniaceae were by far the most abundant microeukaryote throughout the experiment (Supplementary Figure 4). Outside of Symbiodiniaceae, the protist community under control (C0) conditions was dominated by *Prymnesium* (Haptophyta) and *Rhodomonas* (Cryptophyta) (Figure 1A, Figure 2). Both C0 and HS samples showed internally consistent protist community composition within their respective treatments (Figures 1-2), but HS samples diverged from this baseline through increased dominance of corallicolids, a shift that coincided with their separation from other treatments in the Aitchison distance PCA (Figure 1B & Figure 2). By contrast, MS and CS samples showed greater within-group variability in protist composition, suggestive of a transitional microbiome under abrupt stress, which is consistent with the Shannon Alpha Diversity results (Figure 1A). This was especially pronounced in the CS group, which included a strong outlier suggesting a particularly distinct protist community in at least one cold-shocked coral (Figure 1B).

Several chlorophyte-related genera such as *Ulothrichales* (Chlorophyta_X) and *Cladophora* (Chlorophyta_X) were more abundant in CS and MS samples, before appearing at low abundance in the HS samples. In comparison, corallicolids remained stable throughout most of the experimental thermal stress before significantly increasing in abundance at the HS timepoint (Figure 2). As thermal stress increased, corallicolids showed a significant increase in abundance as early as the mild stress timepoint (MS) compared to C0 (Figure 3; ANCOM-BC, LFC = + 1.29, P=0.00). This increase was even more pronounced by the HS timepoint compared to C0 (Figure 3; ANCOM-BC, LFC = + 2.56, P=0.00), highlighting its growing dominance during heat stress. In contrast, *Rhodomonas*, a photosynthetic cryptophyte, showed substantial depletion in both MS (Table 1; ANCOM-BC; LFC = -1.650, P=0.00) and HS (Table 1; ANCOM-BC, LFC = -4.315, P=0.00) samples. Interestingly, *Symbiodinium* increases in MS (Table 1; ANCOM-BC, LFC = +1.974, P=0.00), but then decreases in relative abundance at the HS timepoint (Table 1; ANCOM-BC, LFC = -0.708, P=0.00). Overall, these patterns indicated a shift from a stable protist community toward one increasingly dominated by corallicolids as thermal stress intensifies.

**FIGURE 3.**
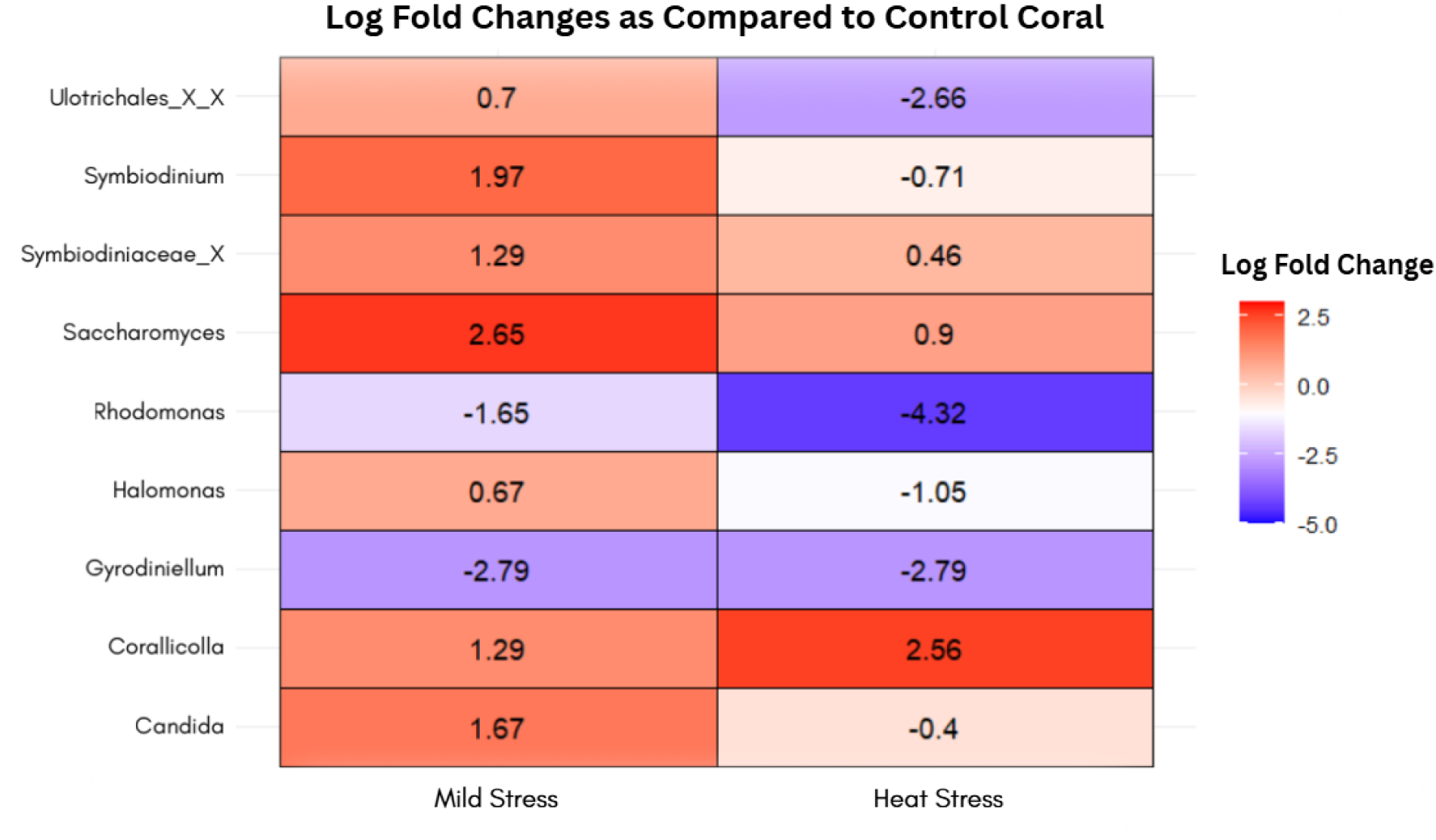
Genus-level heat map shows log-fold changes in protist relative abundance under mild (MS) and severe heat stress (HS) compared to control corals (C0). The Genera-level heat map was generated with ANCOM-BC2 & ggplot2, showing log fold changes in protist relative abundance under mild and severe heat stress relative to C0. Positive values (red) indicate enrichment under mild and heat stress, and negative values (blue) indicate depletion throughout mild and heat stress. Color intensity corresponds to the magnitude of the log fold change, highlighting taxa with the greatest shifts in abundance across treatments.

**TABLE 1.**
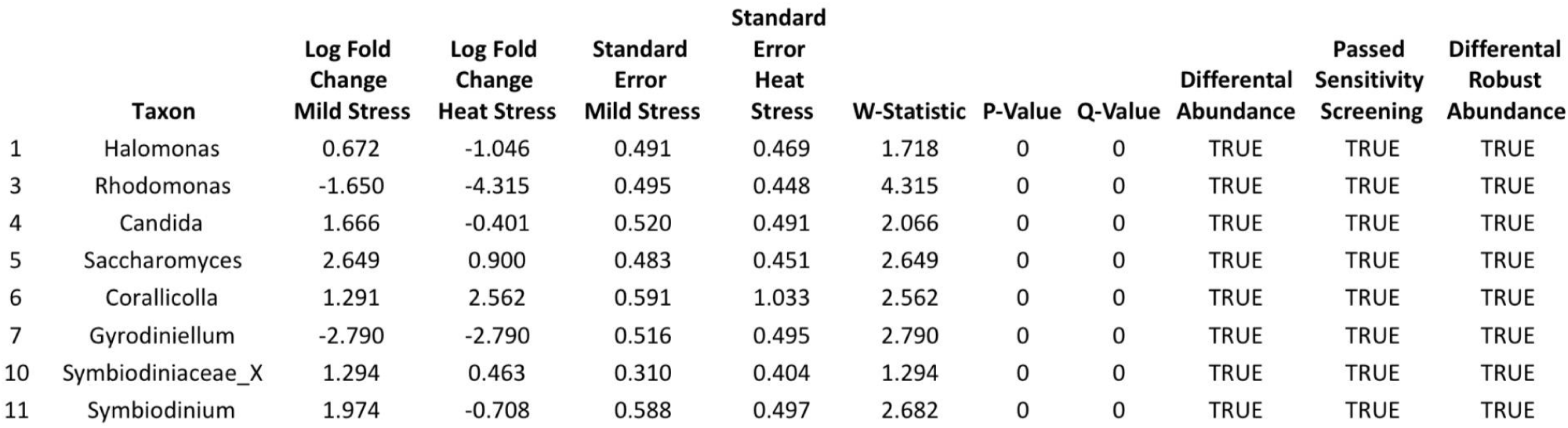
ANCOM-BC significance chart. This chart shows protist genera whose changes in relative abundance across thermal stress conditions was significant (q-value <= 0.05). Corallicolida showed a significant increase in abundance under heat stress indicating that this increase was consistent across samples and not attributable to random variation.

## DISCUSSION

Previous research has shown that *Pocillopora* corals generally maintain a stable prokaryotic microbiome under stress (Pogoreutz et al. 2018; Ziegler et al. 2019; Bergman et al. 2021); however, the protist portion of its microbiome has remained unexamined in this context. Using anti-metazoan 18S rRNA gene metabarcoding, we show that the protist-associated microbiome (eukaryome) of *P. damicornis* exhibits a distinct response to heat stress. Specifically, we see corallicolids increase dramatically in relative abundance during heat stress while many other protist taxa declined. This pattern is consistent with previous reports linking a higher corallicolid abundance to thermally susceptible soft corals and eukaryotic destabilization during bleaching events in stony corals (Bonacolta et al. 2024; Peterson et al. 2025). The significant enrichment confirmed by ANCOM-BC further demonstrates that this pattern most likely represents a consistent biological response rather than random variation. Corallicolids remain understudied with respect to their ecological role and effect on host fitness. They are known to live intracellularly within coral mesenterial filaments and have not been associated with overt host pathology under stable environmental conditions (Kwong et al. 2019). The dramatic increase observed during heat stress suggests that this relationship may become opportunistic as the coral host is weakened by external stressors. Corallicolids may capitalize on the dysbiotic conditions underlying coral bleaching and the weakened immune system of the coral host, allowing them to thrive while other members of the microbiome decline. These patterns support the hypothesis that corallicolids may transition from a largely commensal role toward a potentially parasitic role under unfavorable environmental conditions for the coral.

While corallicolids became dominant throughout thermal stress, many other protist taxa declined substantially. *Rhodomonas*, a photosynthetic cryptophyte that can serve as a heterotrophic food source for corals (Leal et al. 2014), declined during both mild and severe heat stress. Experimental evidence has shown that several coral species selectively capture this microalgae (Leal et al. 2014), *Rhodomonas marina*, indicating that the decrease in this cryptophyte may represent a decrease in the available supplemental nutrition during periods of thermal stress. Likewise, several chlorophytes, including *Ulothrichales* and *Cladophora*, became less abundant under heat stress, suggesting that *Pocillopora*-associated green algae are particularly sensitive to elevated temperatures. Symbiodinium displayed a different response, increasing during mild thermal stress before significantly declining during severe heat stress. This initial spike likely reflects the coral host losing control of its algal symbiont population, due to the destabilized symbiosis (Rädecker et al. 2022). The shift toward a eukaryome increasingly dominated by corallicolids likely reflects dysbiotic conditions and may further compromise the coral’s ability to withstand environmental stress (Marangon et al. 2025). As corallicolid abundance increases, the coral holobiont may become increasingly vulnerable to additional environmental stressors as the host contends with this higher parasite load.

Managing the coral tank allowed direct control and monitoring of all systems including temperature, salinity, nutrient availability, alkalinity, and lighting. This design enabled close observation of coral stress progression but also introduced variability due to the small-scale, manually regulated aquarium system. These uncertainties should be considered when interpreting changes in microbiome composition. A distinctive part of this experiment was the inclusion of an unplanned cold stress event. While attempting to reduce the rate of temperature increase during the heating treatment, after rapid bleaching was observed, a cooling fan was introduced and left on long enough to cause a substantial drop in temperature below 18℃. This resulted in a cold shock to the coral fragments. Rather than excluding this event, samples were collected to explore how sudden cold stress might influence the protist microbiome. While the microbiome was more diverse during the cold stress event compared to the control, the two most abundant taxa outside the Symbiodiniaceae were *Corallicolida* (Apicomplexa) and *Pseudocolliniidae* (Ciliophora), a family of heterotrophic ciliates containing several known parasites of marine invertebrates (Chantangsi et al. 2013). Although there is no clear known role in the coral microbiome, their increased abundance may reflect a shift in the protist community towards opportunistic pathogens during cold stress. Cold stress in corals has been studied far less extensively than heat stress, yet it may disrupt host function and symbiotic communities in a similar way. It is therefore possible that cold stress may induce microbiome changes comparable to those observed during heat-induced bleaching.

While this study provides valuable preliminary results, the modest sample size limits the generalizability of the findings. Nonetheless, this study provides a framework for examining protist-driven microbiome responses across multiple thermal stress conditions within a controlled system. By testing multiple stress states, including an accidental but ecologically relevant cold shock, this work expands our knowledge of how coral-associated protists respond to thermal stress.

## CONCLUSION

Experimental thermal stress caused significant shifts in the eukaryome of *P. damicornis,* marked by increased dominance of corallicolids (Apicomplexa) during heat stress. ANCOM-BC identified significant enrichment of corallicolids under heat stress, indicating this shift likely reflects a consistent, stress-associated response rather than random variation. These microbiome changes occurred alongside visible bleaching and decline in coral health, consistent with the hypothesis that thermal stress disrupts the normally stable coral-corallicolid relationship. These findings align with, and extend, prior work linking corallicolids to thermally susceptible corals by demonstrating corallicolid enrichment across multiple thermal stress states. While corallicolids persist at low, apparently benign levels under stable conditions, their pronounced increase during heat stress raises the possibility that they shift toward a more opportunistic or parasitic role as host health declines. Together, these results position corallicolids as a key indicator of protist-associated microbiome instability during thermal stress, underscoring the importance of understanding protists’ role in the coral microbiome for predicting and addressing coral vulnerability under climate change-driven warming.

## ACKNOWLEDGEMENTS

We would like to thank Dr. Leocardio Blanco Bercial and Dr. Liv Williamson for putting the authors in contact. This study was supported by funds from the Steamboat Springs High School, donations from Underwater Wonders (Fort Collins, CO), and a grant from the DonorChoose Program funded by Colorado Gov. Jared Polis’ administration. A.M.B. is supported by a Postdoctoral Fellowship from the Natural Sciences and Engineering Research Council of Canada (NSERC; CPRA - 608874 – 2026).

## Data Availability Statement

Raw sequencing data has been deposited onto NCBI SRA under BioProject PRJNA1475488. Code used for sequence analysis, as well as ASV sequences, count table, and taxonomy table can be found on GitHub at: https://github.com/CoralCode1/Pdam_corallicolids/

## SUPPLEMENTARY INFORMATION

**Supplementary Figure 1.**
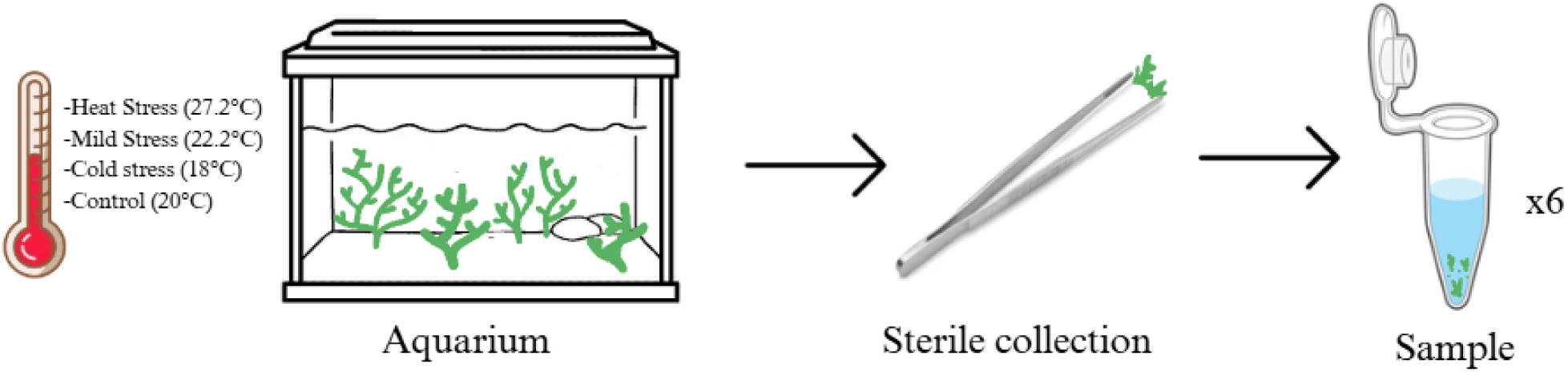
Experimental methods. The sample collection is shown from the temperature variance in the tank, the coral living in the tank, and the randomized sterile collection of the polyps which were placed in tubes, repeated 6 times for each temperature time point.

**Supplementary Figure 2.**
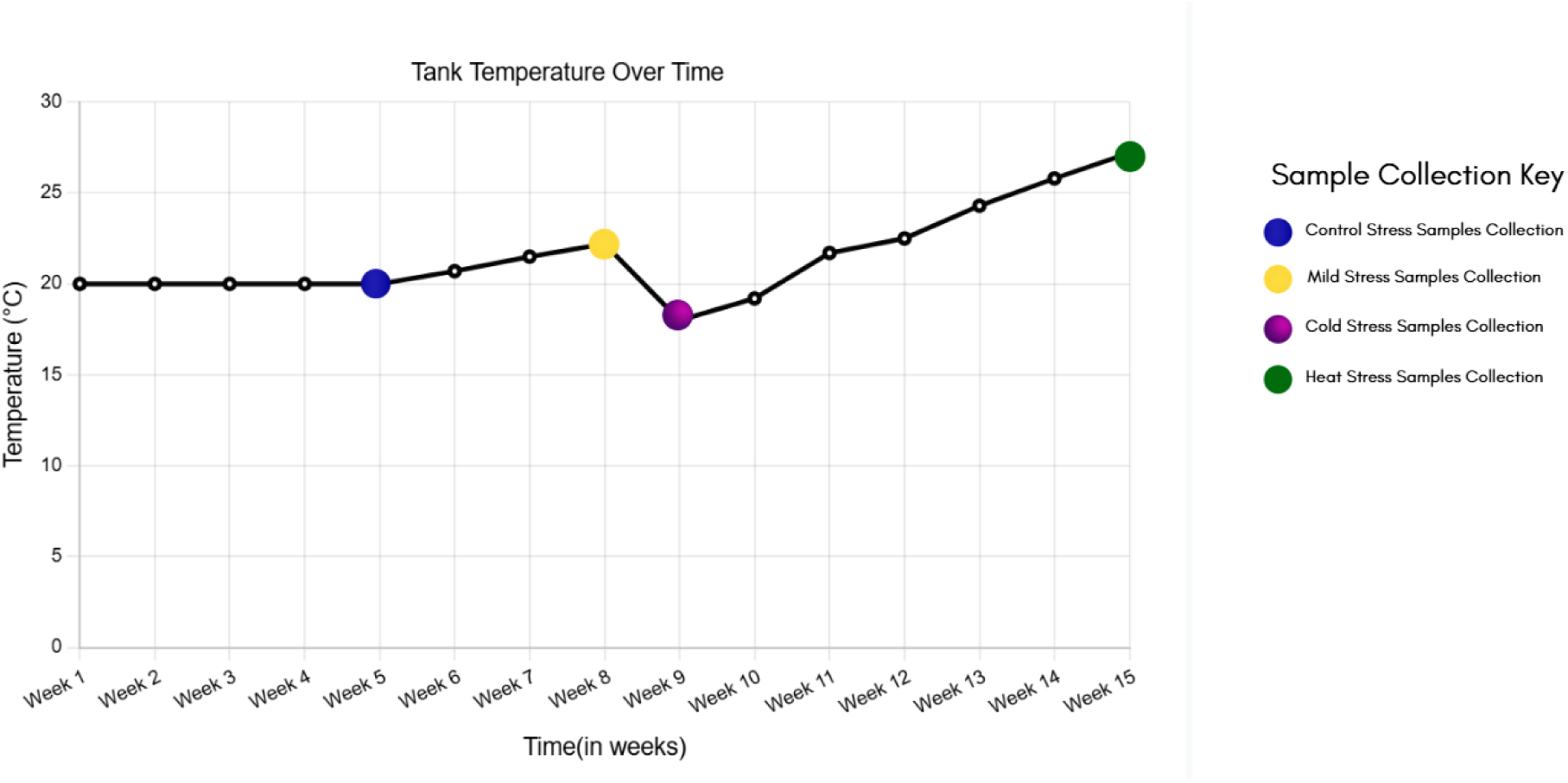
Line plot demonstrating tank temperature throughout experiment. The average tank temperature of each week is shown through the progression of the experiment. Colored dots indicate when sampling was performed.

**Supplementary Figure 3.**
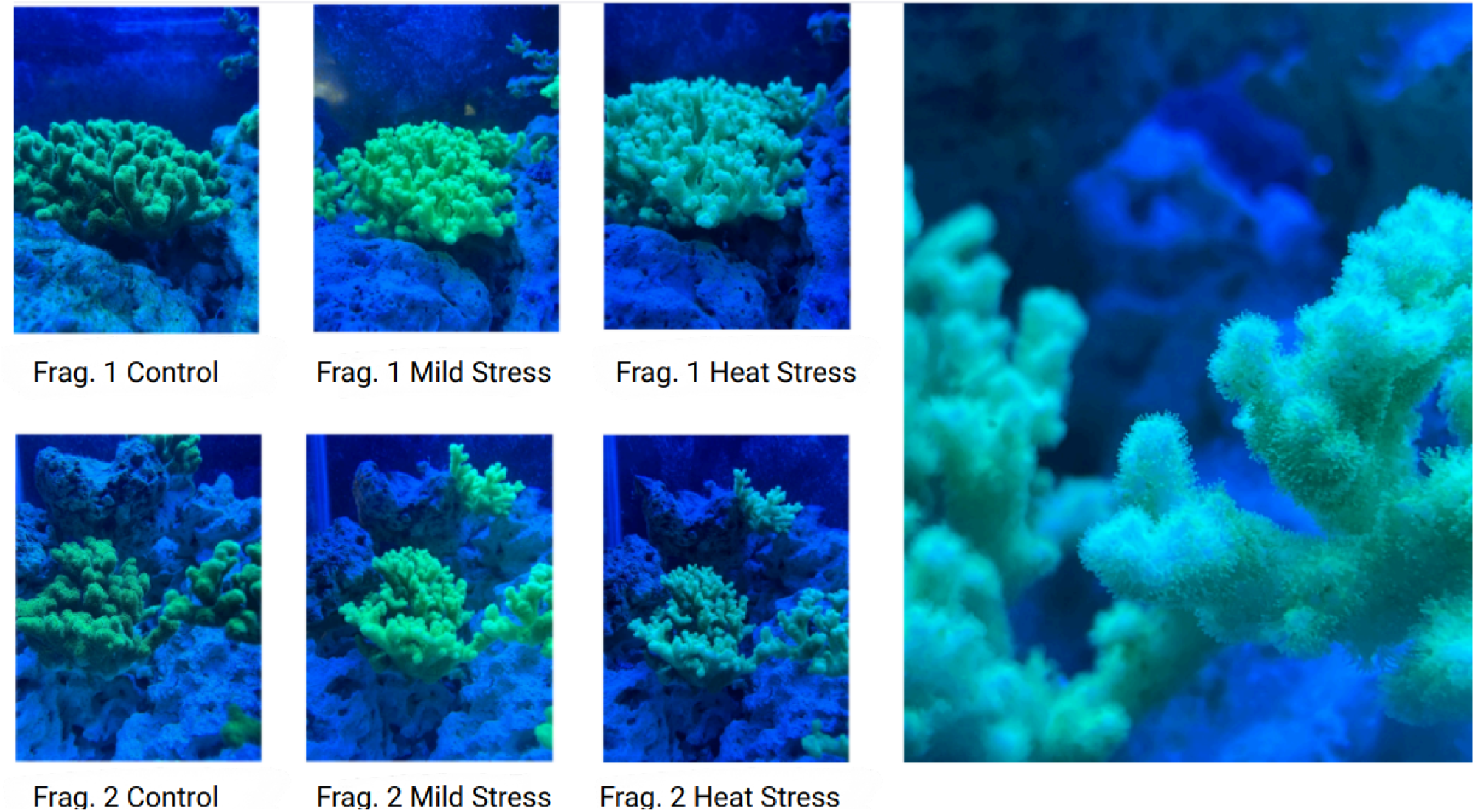
Coral fragments throughout thermal stress. Images of the coral fragments tested in the experiment show the visible effect of thermal stress throughout the different stress events. The coral begins in a healthy, dark green state (Control, C), becomes fluorescent, a key stage in the bleaching process as the coral brightens as a stress response (Mild Stress, MS), and whitens as the symbionts are expelled from the coral (Heat Stress, HS). No pictures of the cold stress were recorded.

**Supplementary Figure 4.**
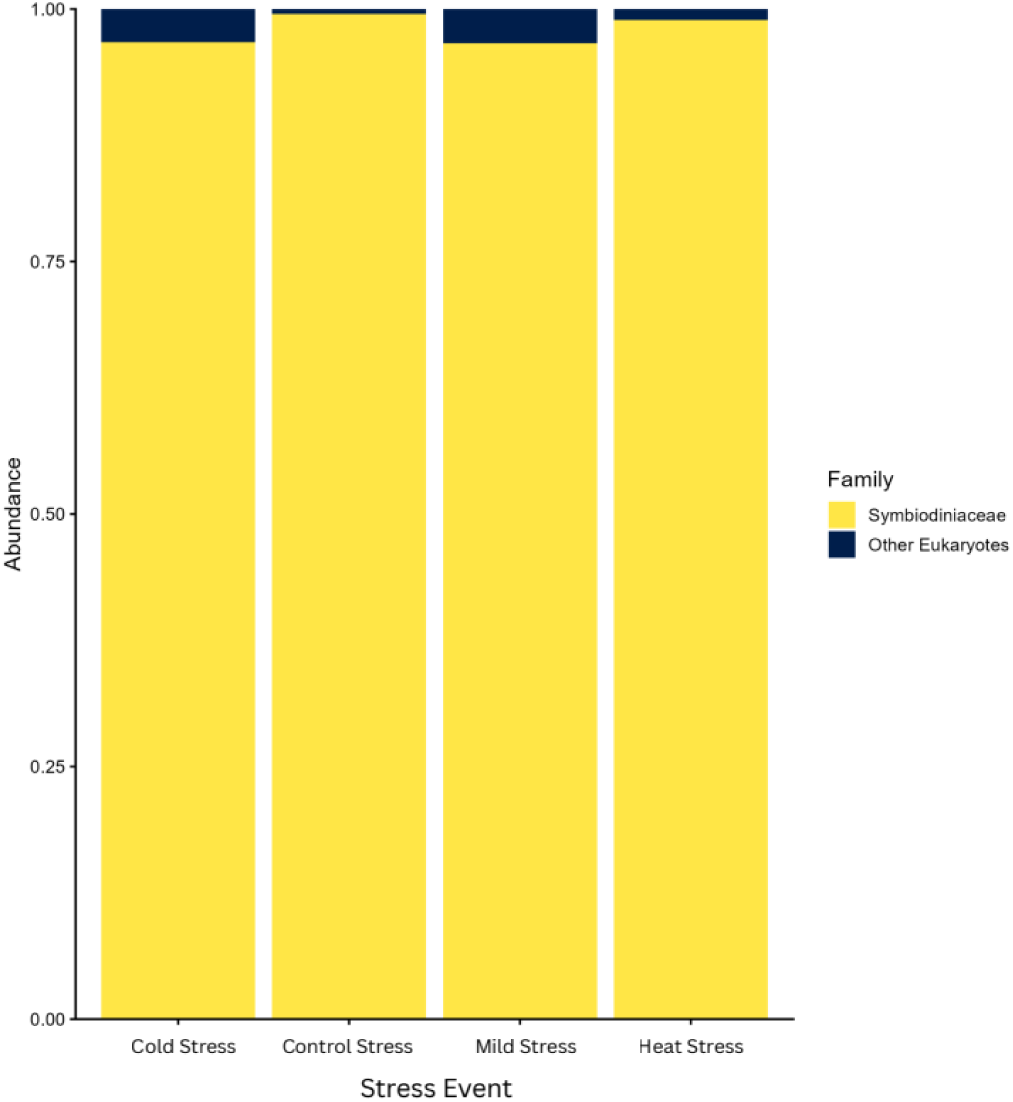
Symbiodiniacae Plot. Bar plot of Symbiodiniceae relative abundance through the experiment.

